# Privacy-Preserving Genotype Imputation in a Trusted Execution Environment

**DOI:** 10.1101/2021.02.02.429428

**Authors:** Natnatee Dokmai, Can Kockan, Kaiyuan Zhu, XiaoFeng Wang, S. Cenk Sahinalp, Hyunghoon Cho

## Abstract

Genotype imputation is an essential tool in genetics research, whereby missing genotypes are inferred based on a panel of reference genomes to enhance the power of downstream analyses. Recently, public imputation servers have been developed to allow researchers to leverage increasingly large-scale and diverse genetic data repositories for imputation. However, privacy concerns associated with uploading one’s genetic data to a third-party server greatly limit the utility of these services. In this paper, we introduce a practical, secure hardware-based solution for a privacy-preserving imputation service, which keeps the input genomes private from the service provider by processing the data only within a Trusted Execution Environment (TEE) offered by the Intel SGX technology. Our solution features SMac, an efficient, side-channel-resilient imputation algorithm designed for Intel SGX, which employs the hidden Markov model (HMM)-based imputation strategy also utilized by a state-of-the-art imputation software Minimac. SMac achieves imputation accuracies virtually identical to those of Minimac and provides protection against known attacks on SGX while maintaining scalability to large datasets. We additionally show the necessity of our strategies for mitigating side-channel risks by identifying vulnerabilities in existing imputation software and controlling their information exposure. Overall, our work provides a guideline for practical and secure implementation of genetic analysis tools in SGX, representing a step toward privacy-preserving analysis services that can facilitate data sharing and accelerate genetics research.^†^

**Availability:** Our software is available at https://github.com/ndokmai/sgx-genotype-imputation.

## 1 Introduction

While advances in high-throughput sequencing technology have resulted in a large-scale collection of genomic sequences, the cost of sequencing remains prohibitively high for researchers studying large cohorts. As a result, many large-scale studies use array-based or targeted sequencing platforms that characterize only a subset of potential genetic variants [1–3]. Furthermore, even when the data from the whole genome is available, lack of coverage or repetitive sequences can render variant calling inconclusive in key genomic regions [4].

Genotype imputation methods [5–12], which infer missing genotypes in a dataset using a reference panel of high-quality haplotypes, have thus become an essential tool for improving downstream analyses like genome-wide association studies and fine-mapping by amplifying the resolution of the dataset. A number of genomic data repositories now provide genotype imputation as an online service (e.g. Michigan Imputation Server [11]), allowing researchers to impute their data against a large reference panel for improved imputation accuracy. Unfortunately, using these services require the researcher to upload their dataset to an external server operated by the imputation service provider, which raises important security and privacy concerns. To ensure that these services are broadly applicable to a wide range of researchers around the world with varying privacy needs, we require new tools for outsourcing genotype imputation that do not put the privacy of the user’s data at risk.

To this end, we develop a solution based on the rapidly-advancing Trusted Execution Environment (TEE) technologies, in particular the Intel SGX (Software Guard Extensions) framework [13], which is widely available in recent generations of Intel CPUs. Intuitively, SGX provides runtime protection for applications running inside the SGX enclave through page-level memory encryption and access control enforced on a hardware-level. SGX technology has been attracting growing interests from the bioinformatics community [14–16] due to its potential to enable privacy-preserving analysis pipelines for sensitive genetic datasets. In contrast to alternative cryptographic frameworks for secure computation such as homomorphic encryption (HE) [17], the key benefit of SGX is that it incurs relatively small computational overhead, since most of its computation is performed inside the SGX enclave based on the plaintext data. Notably, the recent iDASH-2019 competition Track-2 (http://www.humangenomeprivacy.org/2019/) explored HE-based solutions for privacy-preserving genotype imputation. While the competition led to promising solutions [18, 19], existing proposals are likely to face additional challenges for adoption due to their reliance on simplified imputation algorithms with limited accuracy, as we show in our results.

Despite the promises of Intel SGX technology, deploying a fast and secure program in SGX is a highly non-trivial task for a number of reasons. First, because the security of Intel SGX relies on assumptions about the hardware environment, there are a range of potential attack vectors (e.g. side-channel attacks due to memory access patterns, runtime differences, or resource contention [20–22]) that need to be taken into consideration when developing SGX-based solutions. As we demonstrate in our work through a range of example attacks, off-the-shelf deployment of existing software to the SGX enclave provides only partial protection for the data, which may not meet the security needs for tasks involving sensitive genetic information. Furthermore, the SGX architecture increases the cost of memory access due to an additional layer of encryption introduced by SGX. As a result, programs need to be carefully designed to optimize memory and cache usage to avoid excessive overhead in runtime.

In this work, we overcome these challenges to introduce SMac, a scalable and privacy-preserving genotype imputation algorithm for Intel SGX. Our imputation strategy closely follows that of Minimac [11], a state-of-the-art imputation tool adopted by the Michigan Imputation Server, while additionally providing secure enclave-based protection for the user’s data. We address the aforementioned pitfalls of SGX by leveraging a range of techniques, including algorithmic design and software implementation strategies to provide comprehensive protection against key side-channel vulnerabilities of SGX and an efficient implementation of the algorithm to minimize memory usage. We evaluate SMac on two real-world datasets, including 1000 Genomes Phase 3 (1KG) [23] and Haplotype Reference Consortium (HRC) [24], and show that SMac achieves the same imputation accuracy as Minimac (in particular Minimac3, the most accurate but slower version of Minimac without the additional heuristics introduced in Minimac4) while additionally preserving the privacy of the user. We also demonstrate that SMac offers significantly better accuracy compared to recent HE-based solutions for genotype imputation. Even with the additional overhead of the SGX environment and the limited RAM available, the computational performance of SMac remains practical, incurring roughly over 50% overhead in runtime compared to Minimac (more specifically Minimac4, the most recent and fastest version of Minimac, which, due to some heuristic shortcuts it employs, is potentially less accurate than Minimac3) on the full HRC dataset including 54k haplotypes. These results demonstrate the effectiveness of SGX-based genotype imputation enabled by SMac. Our open-source implementation of SMac is publicly available at https://github.com/ndokmai/sgx-genotype-imputation

## 2 SMac: Our SGX Solution for Secure Genotype Imputation

Here we describe SMac, our solution for privacy-preserving genotype imputation based on Intel SGX. The overall workflow of SMac (illustrated in Figure 1) is as follows. First, the user sends their encrypted input data (i.e., partially observed genotype profiles to be imputed) to the imputation service provider (SP) equipped with CPUs with Intel SGX support. The integrity of the SP’s CPU and the program binary to run inside the secure enclave (i.e., SMac executable) are verified through the process known as ‘remote attestation’ [25]. Note that SP also holds a “reference panel” (a collection of fully-sequenced haplotypes, see the definition below) to use for imputation, which is additionally provided as input to SMac; since SMac runs on the SP’s system, the reference panel is considered public data for our purposes, although this is not a strict requirement for the feasibility of SMac. Once remote attestation is successfully completed and the user input received by the SP, SMac runs in the SGX enclave to impute the user’s data. The encrypted user input as well as any intermediate data (in RAM) used by the SGX process are decrypted only within the enclave, a security property that is enforced at the hardware level. Importantly, SMac employs additional techniques we introduce to provide protection against key side-channel leakages of sensitive information, thus providing stronger security properties than conventional approaches to deploying an SGX application. Once the whole data is imputed by SMac, the imputed genotypes are returned to the user in an encrypted form, to be locally decrypted by the user to obtain the final results. The implementation details of our system architecture are included in the Appendix. In the following sections, we describe our design strategies for SMac in more details.

**Figure 1:**
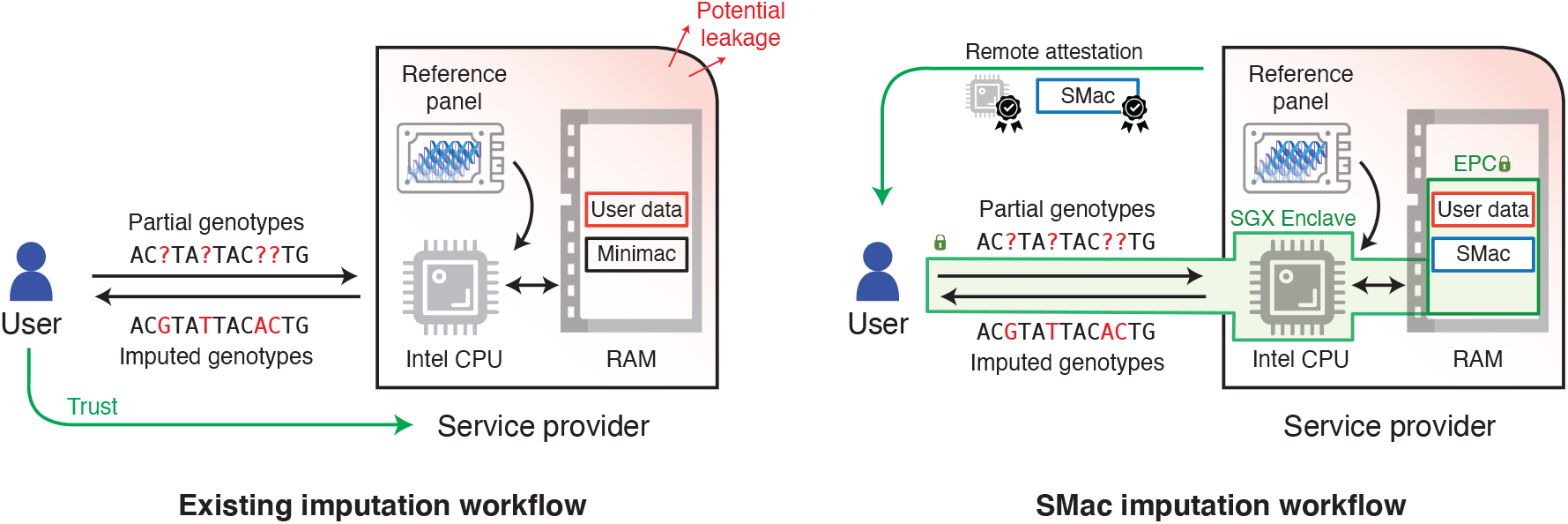
Overview of SMac. A comparison of the SMac workflow (right) with that of existing imputation servers based on Minimac (e.g. Michigan Imputation Server) (left). Key differences include remote attestation (which verifies the integrity of the hardware/software configuration of the server to the user), end-to-end encryption of input (partial) and output (imputed) genotype profiles, and processing the reference panel as a stream due to SMac having access to only a limited portion of RAM (EPC) in the SGX enclave. SMac enables genotype imputation service with stronger security guarantees. CPU: central processing unit. RAM: random access memory. EPC: enclave page cache.

### 2.1 Overview of Genotype Imputation

We start by providing a high-level description of the genotype imputation algorithm implemented in SMac, which employs a hidden Markov model (HMM)-based approach introduced by Minimac [11, 26], a state-of-the-art software tool for genotype imputation (adopted by Michigan Imputation Server). The input to SMac includes: (i) a *target haplotype* from the user, represented as a sequence of *n* genotypes 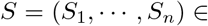 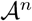, a subset of which are unobserved latent variables, where 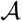 denotes the alphabet (e.g. 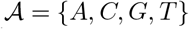 and (ii) a *reference panel*, which consists of *m* fully-observed haplotypes 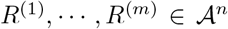 to be used for imputing the user’s data. Let *O ⊂* [*n*] be the subset of indices where *S*_*i*_ ∈ O is observed in the target haplotype, and define *S*_*O*_ ≔ (*S*_*i*′∈*O*_)_*i′∈O*_. Let *M* = [*n*] *\ O* be the set of missing genotypes in the target haplotype to be imputed. The goal of SMac is to infer the missing genotype *S*_*i*_ for each *i ∈ M*, based on the input reference panel and observed genotypes *S*_*O*_. Specifically, we want to compute the conditional distribution ℙ(*S*_*i*_|*S*_*O*_) for each missing genotype *i ∈ M*. Leveraging the HMM as a model of choice for representing the genotype distribution ℙ(*S*_1_*, …, S*_*n*_), a classical and widely-used technique in genetics [27], the computation of beliefs over the missing genotypes can be accomplished using a straightforward dynamic programming algorithm [28] (also known as the forward-backward algorithm) with *O*(*mn*) running time to impute the target haplotype. The running time can be significantly reduced in practice by exploiting the similarities among haplotypes in small genomic segments [11, 26, 29–34]. Following the approach of Minimac, SMac takes advantage of this redundancy by taking as input a compressed reference panel that has been partitioned into consecutive *blocks* with distinct haplotypes within each block combined. We include the details of SMac imputation algorithm in the Appendix.

### 2.2 Overview of Intel SGX Security Model and Its Limitations

Intel SGX technology [35] offers hardware-based protection for applications running inside an SGX enclave against a potentially malicious kernel or operating system. The security claims of SGX include the confidentiality and integrity of data and application binary at runtime. These are achieved via hardware-level cryptographic building blocks, access control mechanisms, and authenticated encryption engine for memory pages built into all Intel CPUs since the 6th generation (released in 2015). In the SGX environment, data is always kept encrypted and only decrypted within the CPU itself for processing. In comparison to software-based protection (e.g. virtualization), compromising hardware-based security requires more sophisticated attacks against the complex hardware architecture and sometimes the physical hardware itself.

A common use case of SGX is for secure *outsourced* computation (e.g. a genotype imputation service), where a user sends private data to a remote SGX enclave, controlled by an untrusted party, for secure data processing. To achieve desired security properties in this remote setting, Intel SGX relies on a process known as *remote attestation*. By taking advantage of a hardware-level digital signature component, remote attestation provides a proof of authenticity to the user, which ensures the integrity of the application binary running inside the SGX enclave as well as the security of the communication channel between the user and the remote enclave (thus avoiding man-in-the-middle attacks).

Despite the security claims of SGX, there are additional security challenges that practitioners looking to leverage this technology need to take into consideration. First, Intel takes note of four *side-channel* attack vectors (i.e. leakage of sensitive information through monitoring of computer system behaviors external to the program itself) that are not included in the SGX security model: power statistics, cache miss statistics, branch timing, and page accesses via page tables [13, 36]. Understandably, this design decision is aimed at reducing the complexity of the architecture and thus the performance penalty and attack vectors. However, this implies that additional mitigation strategies need to be employed at the software level in order to achieve strong security properties. Our work aims to address this limitation of SGX by devising algorithmic design and implementation strategies to protect against key side-channel vulnerabilities in the SMac threat model (described in the next section).

We note additional hardware vulnerabilities and attacks related to SGX have been discovered by researchers in recent years. These include L1 Terminal Fault [37, 38], also known as Foreshadow, Microarchitectural Data Sampling [39–42], and power side-channel attacks in PLATYPUS [43]. To mitigate these attacks, Intel has recently published microcode updates and added silicon-based protection to new Intel CPUs [44–46]. We are able to verify that Microsoft Azure Confidential Computing [47] offers SGX cloud servers with all the mitigation in place by default. We recommend practitioners who wish to deploy SGX applications to strictly follow mitigation guidelines or use SGX cloud services that take serious precaution against these vulnerabilities.

### 2.3 Our Threat Model and Attack Vectors

In our threat model for SMac, the service provider (SP) is assumed to be malicious and has complete control over the operating system, the kernel, and I/O of running enclaves. In other words, the SP is an *active* adversary who may attempt to eavesdrop and/or tamper with any steps of SGX enclaves and communication channels in and out of them. The goal of the malicious SP is to extract the genotype data belonging to the user, potentially through intermediate results of SMac computation that may indirectly reveal the underlying genotypes. We assume that the SP has full knowledge of the reference panel.

We rely on the standard SGX security model as previously discussed in Section 2.2, where memory access patterns and timing information are considered attack vectors. We thus assume that known hardware vulnerabilities have been properly mitigated according to Intel’s guidelines [44, 45]. With the SGX remote attestation protocol in place, we assume the existence of a secure authenticated channel between the user and an SGX enclave and that the SMac binary running inside the enclave is authentic and verified by the user. We exclude physical attacks (e.g. bus tapping, power analysis) from consideration due to the difficulty and high cost of performing (and mitigating) this class of attacks.

To focus our efforts on addressing realistic attack scenarios for genotype imputation in SGX, we carefully reviewed the Minimac algorithm in the context of known SGX side-channel attack vectors in the literature. We discovered *two* main attack vectors, described below. Note that we demonstrate in our results how one could exploit these vulnerabilities to extract user genotypes when these threats are not properly mitigated.

#### (i) Secret-dependent timing differences in floating point operations

Andrysco *et al.* [20] have demonstrated that floating point arithmetics processed by Intel CPU’s floating point unit (FPU) can result in a variety of timing statistics based on the provided input values. This timing discrepancy is most pronounced between normal and subnormal (i.e. small values near zero) floating point operations (a relative difference of more than 100% for multiplication and division). Interestingly, this discrepancy is the result of FPU optimization. Although such an optimization is immensely useful in many applications, for a security-critical task like genetic data analyses, it unfortunately also exposes an attack vector. While the user’s genotypes in Minimac are not directly encoded as floating points, various branching decisions based on the user input can lead to different classes of floating-point operations being performed, thus generating timing statistics that can leak the underlying genotypes.

#### (ii) Secret-dependent memory access patterns

We found that Minimac exhibits both *spatial* and *temporal* memory access patterns dependent on the user’s genotypes in L1, L2, last-level cache (LLC). Spatial memory access patterns are exposed by branching decisions based on the user’s genotypes, whereby the two branches access two different memory addresses. Temporal access patterns are exposed indirectly when the temporal gap between two instances of memory access calls depends upon the secret due to timing differences in the instructions carried out inbetween (e.g. timing discrepancies in floating point operations). As our results show, all of these attack vectors present a real threat for the security of SGX-based genotype imputation, thus motivating our mitigation strategies introduced in the next section.

### 2.4 Our Side-Channel-Resilient Implementation Techniques for SMac

To prevent attacks against SMac leveraging side-channels including the ones presented in the previous section, we have developed and employed key algorithmic and software design techniques in the implementation of SMac to provide security against side-channels while preserving the accuracy of the imputation results. Our main strategy is to adopt leakage-resilient arithmetics and branching operations to replace all vulnerable codes in SMac, leveraging CPU instructions that are known to run in constant-time regardless of the input. In order to solely rely on leakage-resilient operations while enabling accurate calculations, we transformed the core imputation algorithm used by Minimac to use fixed-point integers to represent continuous values in the log-domain. This required us to develop novel leakage-resilient subroutines for key operations in our algorithm, including log-sum-exp (LSE) and log-diff-exp (LDE) functions. Lastly, we devised a strong typing-based programming framework to enforce leakage-resilience of the overall SMac program *at the syntactic level*. We describe each of these approaches in more detail below.

#### Constant-time CPU instructions

We used constant-time CPU instructions as building blocks to implement leakage-resilient arithmetics and branching operations to replace all vulnerable codes in SMac. Specifically, we follow Andrysco *et al.*’s argument [20] that floating-point arithmetics (e.g. `fadd, fsub, fmul, fdiv`) and integer division (e.g. `idiv`) instructions on Intel CPU can expose a timing side-channel, while other integer operations (addition, subtraction, multiplication, comparison, and bit-shifts) and Boolean operations run in constant-time. We empirically validated these well-established claims in our experimental setting (data not included). We therefore set out to design our leak-resilient operations around these constant-time instructions. Note that branching operations in the program can be safeguarded by evaluating both branches and selecting the intended output via a multiplexer (i.e. multiply with a one-hot vector of binary indicators across the branches and sum the results). In the following, we describe how all aspects of the imputation algorithm can be implemented using only this limited set of safe operations.

#### Fixed-point representation

First, we sought to eliminate floating-point instructions entirely (given the timing leakage in floating-point operations we previously described) by representing real numbers as *fixed point* integers. Intuitively, fixed point representation maps a real number to an integer by scaling it up by a constant factor. For example, 1.39485 can be represented as ⌊1.19485 × 2^20^⌋. Addition and subtraction of numbers in the fixed-point representation follow naturally from the underlying integer arithmetics as long as the scaling factors match. Multiplication, however, will cause a shift in the scale, e.g. ⌊8.28 × 2^20^⌋ × ⌊0.5 × 2^20^⌋ ≈ ⌊4.14 × 2^20^⌋ /2^20^. We append to each multiplication a re-scaling step that applies a bit-shift (also a safe operation) to truncate the least significant bits.

A key hurdle imposed by our reliance on fixed-point representations is that of *precision*. Fixed point values generally require more bits of information than floating points to represent small numbers. In practice, using the native 64-bit integers in the Intel x86-64 to represent fixed points with a reasonable choice of scaling factor of 2^20^ will limit the smallest number that can be represented to 2^*−*20^ ≈ 9.54 × 10^*−*7^. We have found that this is a vastly insufficient level of precision to produce meaningful imputation results. Indeed, an existing general-purpose library for fixed-time fixed-point arithmetic [20] uses 64-bit integers, leading to overwhelming precision loss for our application. In principle, 128- or 256-bit integers can be used instead if they are natively supported by the CPU and are deemed to be safe; however, this approach effectively doubles or quadruples memory usage while still providing limited precision compared to floating point operations.

#### Log-transformed fixed points

To address the above challenge, we opted to *log-transform* the floating points prior to fixed-point conversion for use in our side-channel resilient program. This led to key improvements in the precision of our imputation algorithm; in fact, our experiments demonstrate that fixed-point representations based on *32-bit* integers are sufficient to achieve highly accurate results for genotype imputation. Note that the existing Minimac software do not perform calculations in the log-domain, instead relying on the precision of floating point numbers. Another major benefit of the log-transformation is that it converts pairwise multiplications and divisions in the algorithm to additions and subtractions, respectively, which are efficient and naturally leakage-resilient in the fixed-point setting. We note that the HMM-based imputation algorithm heavily relies on multiplications, as the algorithm computes aggregate products of probabilities for the latent variables as it walks along the genetic sequence; thus, our log-transformation also helped us to achieve improvements in runtime.

#### Leakage-resilient subroutines for log-transformed fixed points: LSE and LDE

A key difficulty in working with log-transformed numbers is that addition and subtraction in the original space becomes *non-linear* in the transformed space, which is more difficult to perform using only the safe operations available to us. We refer to these required subroutines as log-sum-exp (LSE) and log-diff-exp (LDE), which implicitly implement addition and subtraction in the original space, respectively. In theory, one could define leakage-resilient subroutines for log and exp functions (e.g. based on polynomial approximations), then use them to map the numbers to the original space, perform addition or subtraction, then back to log-domain to implement LSE and LDE. However, directly evaluating log and exp is numerically unstable given the high non-linearity of these functions, and the fact that the intermediate results of these steps being represented as fixed points in the original space incurs precision loss and, to an extent, defeats the purpose of our log-transformation strategy. Thus, we instead developed leakage-resilient subroutines that *directly* approximates the LSE and LDE functions based on piecewise polynomial functions, which we found to be more accurate and efficient than the aforementioned naïve approach. We give the details of these subroutines in the Appendix.

#### Strong typing for syntactic enforcement of leakage-resilience

To guarantee that all vulnerable codes in SMac have been replaced, we design a strong timing-protected type to represent user’s genotypes and override all of its unsafe operations with our safe ones. On a syntactic level, this guarantees that it is impossible to accidentally leak user’s genotypes under the standard SGX assumption unless we *explicitly* reveal them. The security of our program is thus reduced to the security of the leakage-resilient elementary operations, which we have thoroughly audited. We include a formal definition of the strong typing system we designed and used for the implementation of SMac in the Appendix.

Taken together, these techniques enabled us to implement the imputation algorithm in SMac in an accurate, efficient, and leakage-resilient manner. We included technical details of the software implementation of SMac as well as further optimization strategies in the Appendix.

## 3 Experimental Results

### Side-channel vulnerabilities of SGX threaten the security of genotype imputation in secure enclaves and motivate our mitigation strategies

To demonstrate the challenges of safely deploying existing genotype imputation pipelines to trusted execution environments, we identified two vulnerable “gadgets” in Minimac that can leak sensitive user genotypes in SGX via side-channels to the adversary controlling the imputation server (Figure 2). The first gadget is a part of the *renormalization* step in Minimac: if intermediate values fall below a certain threshold, Minimac multiplies them with a fixed constant to avoid numerical underflow. Because the sizes of the intermediate values are determined by user genotypes, knowledge of this branching decision can be used to infer them. The second gadget is a part of the *emission* step in Minimac, where the value of emission probability (*e*_1_ or *e*_2_ in the figure) used to update the beliefs over the hidden states in the HMM is determined by whether the user genotype matches the reference genotype at a given position. The timing discrepancy caused by multiplication of two different emission probabilities can expose the user’s genotype.

**Figure 2:**
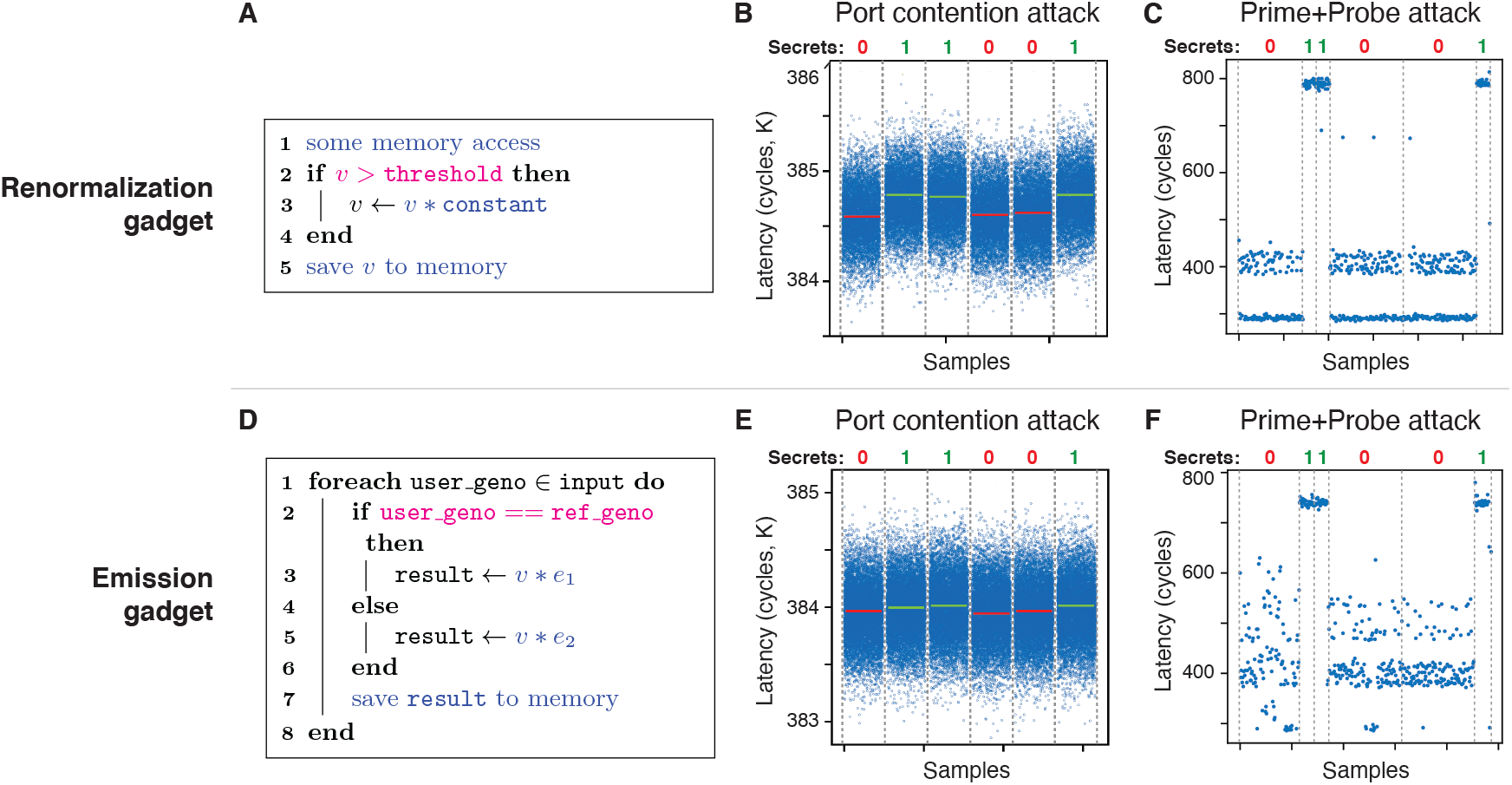
Demonstration of side-channel vulnerabilities in the original Minimac imputation algorithm. We identified two vulnerable gadgets in Minimac—renormalization (top row) and emission (bottom row)—and demonstrated two types of attacks (port contention and Prime+Probe), as described in main text. We provide the pseudocodes of the two gadgets (**A** and **D**) with extractable secrets highlighted in magenta and measurable attack surfaces in blue. **B** and **E** show a successful attack on both gadgets via port contention side-channel. Each data point reflects the amount of contention in clock cycles. Horizontal lines in red and green indicate the sample mean for each segment of the victim process, which successfully distinguish the underlying secrets (0 or 1) shown at the top. **C** and **F** demonstrate another type of attack based on Prime+Probe, which holds even when the simultaneous multithreading (SMT) feature of SGX is turned off as a security measure, unlike port contention. Each data point reflects the latency in the probe step of the attack in clock cycles. Clear distinction in latency is observed between different user secrets (0 or 1) shown at the top.

To demonstrate the vulnerability the two gadgets we identified, we implemented two types of attacks against them based on the port contention [22, 48] and Prime+Probe [36, 49, 50] techniques. Notably, the first attack requires enabling simultaneous multithreading (SMT) in Intel CPU, which is often desirable for performance considerations, while the latter will work even if SMT is disabled as an additional security measure. The attacks are demonstrated in a setting with Intel i7-6700 CPU with Turbo Boost off and in Ubuntu 16.04 and Linux Kernel v4.4.0. Neither of the gadget code runs in SGX during the attacks to simplify memory address discovery, but otherwise we make the same threat assumption as if it were running in SGX. Note that memory address discovery in SGX has been demonstrated in practice [21].

We first performed the **port-contention** attack [22, 48] on a victim process running the gadget code by causing a resource contention at a targeted floating-point unit (FPU). This contention is observed and measured (in the number of clock cycles) to infer how long the gadget code takes to process floating point multiplications, which can leak information about the private input. To boost the signal-to-noise ratio, we apply a recently-proposed MicroScope technique [21] to force the gadget to run repeatedly in speculative execution by triggering page faults at a targeted memory page. We then trained a classifier to distinguish the observed measurement patterns between the two possible user input (0 or 1, representing a genotype) In an experiment with 1,000 randomly sampled bits representing the private input, our attack on the renormalization gadget achieves 99.4% classification accuracy with average runtime of 1.04 seconds/bit, and our attack on the emission gadget achieves the accuracy of 88.5% with average runtime of 1.35 seconds/bit. Example attack results are shown in Figures 2B and 2E. Next, we performed the **Prime+Probe** attack [36, 49, 50] on a victim process running the gadget code by causing cache misses at the last level cache (LLC). The cache-miss statistics of a floating-point multiplication in the gadget code that occur *between* two memory access calls (shown in Figures 2A and 2D) can be measured (in the *probe* step) to infer the duration of the multiplication, and therefore reveal the secret bits. To illustrate that cache-miss statistics can leak the secret information, we modify the gadget to artificially boost the side-channel by making every multiplication repeats itself 1000 times, which in practice could be achieved via techniques such as the MicroScope attack [21]. We show example results based on this attack in Figures 2C and 2F, which demonstrate that there is a clear distinguishable difference between the two possible values for the secret bit. These results provide evidence that side-channel vulnerabilities in Minimac pose a realistic security threat, which we protect against using our constant-time implementation strategies.

### SMac achieves the same imputation accuracy as Minimac while keeping the input data private from the service provider

To evaluate the accuracy of our methods, we used two public datasets commonly used for benchmarking genotype imputation methods: (i) *1000 Genomes Phase 3* (1KG) dataset (5,008 haplotypes from 2,504 subjects) and (ii) *Haplotype Reference Consortium* (HRC) dataset (54,330 haplotypes from 27,165 subjects). We focused our analysis on the genetic variants in the first of the three evenly-sized chunks of chromosome 20 (approximately 23 Mbp in length) generated by Minimac; note that Minimac divides the genome into large overlapping chunks and imputes each chunk separately. This resulted in a total of 401,627 variants for 1KG and 339,328 variants for HRC to be considered in our analysis. We included all types of variants provided in the original datasets, including both single nucleotide polymorphisms and structural variants such as insertions and deletions.

For cross validation, we held out a small subset of subjects for testing (100 subjects for 1KG and 510 subjects for HRC) and used the rest of the dataset as a reference panel to perform imputation on the test data. For the held-out subjects, we considered only the variants on the Illumina Human1M-Duo v3.0 DNA Analysis BeadChip as the observed input data and assessed the accuracy of inferring the rest of the variants in the dataset, reflecting the most common use case of imputation, where the limited data from genotyping arrays are imputed to produce more comprehensive genotype profiles for the study subjects. Notably, the variants on the Illumina array accounted for only a small fraction of the variants in our datasets; 11,012 variants (2.7%) for 1KG and 10,925 variants (3.2%) for HRC were covered by the Illumina array and thus provided to imputation methods as the input. Following the previous work that introduced Minimac, we used squared Pearson correlation coefficients (*r*^2^) between the ground truth and the imputed (expected) allele dosages for the test variants as the evaluation measure, for three different minor allele frequency (MAF) categories: 0.01–0.5%, 0.01–0.5%, and 5–50%.

The results, shown in Figure 3, demonstrate that SMac imputation accuracies are virtually identical to those of Minimac in all MAF categories. In fact, the *r*^2^ value computed for each held-out individual precisely match between SMac and Minimac for all individuals in the test data (Figure 3B). These results illustrate that SMac closely matches the behavior of the state-of-the-art Minimac algorithm, while also providing stronger privacy protection for the user; recall that the SMac imputation pipeline is performed inside the SGX enclave without divulging the user’s input to the imputation service provider. Furthermore, SMac provides comprehensive protection against timing and memory access-based side-channel vulnerabilities, further strengthening its security properties. The fact that there is no loss of accuracy in SMac results is noteworthy considering that SMac employs several approximation techniques to achieve side-channel resilience, including fixed-point arithmetics in log domain and approximations to LSE/LME functions (Section 2.4).

**Figure 3:**
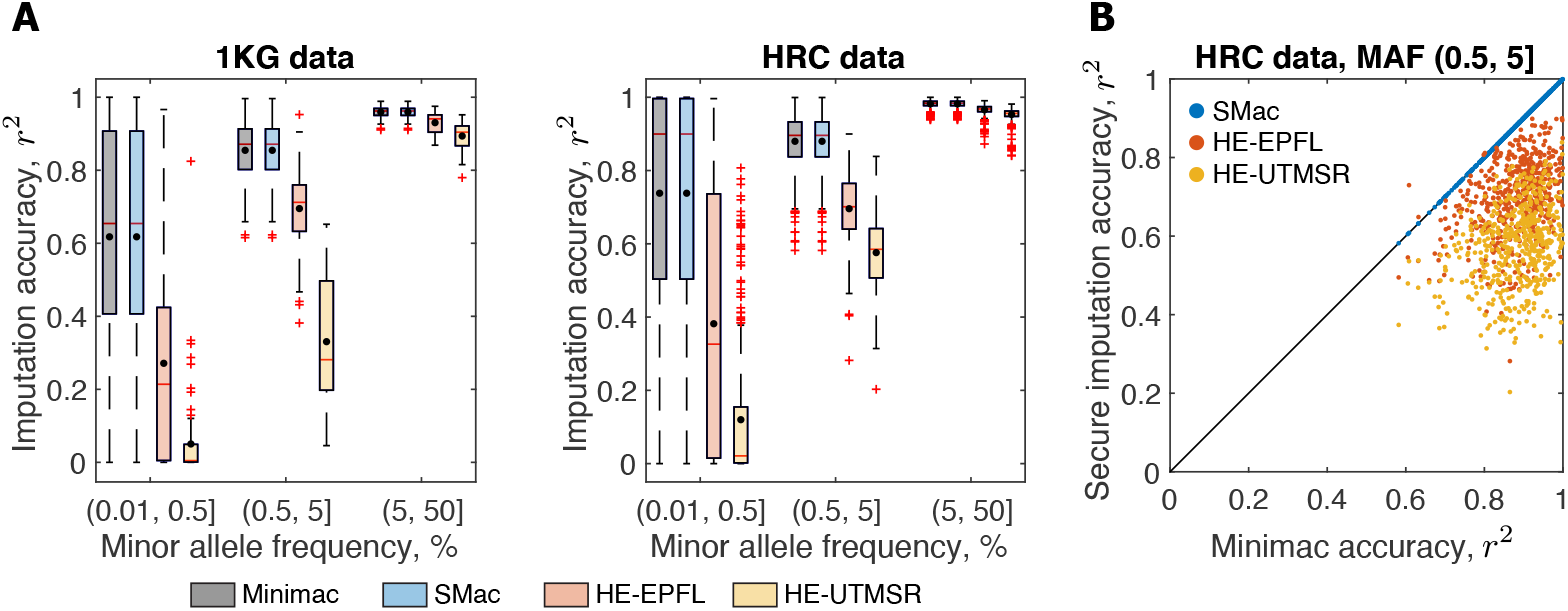
SMac imputation accuracy is identical to Minimac and substantially higher than HE-based solutions. We conducted a cross-validation experiment on 1KG and HRC datasets (chromosome 20) to compare the accuracy of SMac to Minimac4 (the latest version of Minimac), and recent homomorphic encryption-based imputation methods (HE-EPFL and HE-UTMSR) (**A**). Accuracy is measured by Pearson *r*^2^ within each minor allele frequency (MAF) range of the target variant (0.01-0.5%, 0.5-5%, 5-50%). Black dot indicates mean *r*^2^, red horizontal line indicates the median, boxes extend to upper and lower quartiles, and whiskers extend to extreme values excluding outliers marked by red plus symbols. We also plot *r*^2^ of SMac and HE-based solutions for individual test subjects (y-axis) against Minimac *r*^2^ (x-axis) on HRC data, 0.5-5% MAF category (**B**). Note that blue dots representing SMac lie on the diagonal, showing that SMac and Minimac results are identical.

### SMac provides a significant improvement in accuracy compared to homomorphic encryption-based approaches to secure genotype imputation

We additionally compared our method to recently proposed approaches [18] for designing a secure genotype imputation service based on a cryptographic framework known as *homomorphic encryption* (HE). Intuitively, HE is a special form of encryption that allows computation to be implicitly performed over the secrets underlying the ciphertexts by operating only over the ciphertexts (without decrypting them). This powerful technique, in theory, allows arbitrary calculations to be performed over a private dataset without the analyst obtaining any information about the given data, thus promising an enticing solution for secure genotype imputation whereby the user provides data encrypted using HE to the service provider, who then performs imputation and returns the encrypted results to the user without learning any information about the user’s genotypes. However, the key challenge for HE is in developing algorithms that are amenable for efficient computation under HE; certain non-linear operations, such as divisions and comparisons, are costly to perform in HE, often presenting a key bottleneck to achieving practical performance in sophisticated analytic pipelines.

To this end, the iDASH-2019 competition featured a challenge where the participants were asked to develop effective HE-based strategies for secure genotype imputation. Reflective of the aforementioned bottleneck, most of the top solutions employed a *linear* model (e.g. linear or logistic regression) to predict the genotype of a missing variant using the observed variants in the vicinity, representing a considerable departure from the state-of-the-art imputation algorithms such as Minimac. Although these HE-based solutions produced promising performance results [18], we wondered whether our SGX-based strategy could constitute a viable alternative that can provide more accurate imputation performance, notably at the expense of the rigorous privacy guarantee provided by HE, which may be a desirable tradeoff when obtaining accurate results is prioritized.

In our experiments, we evaluated the top two solutions from iDASH based on overall imputation accuracy, which we refer to as HE-EPFL and HE-UTMSR, corresponding to the EPFL (logistic regression-based) and UTMSR (linear regression-based) solutions in the original publication [18], respectively. For the purpose of accuracy comparison, we trained and evaluated the linear models used by the HE-based solutions *in plaintext*, i.e. without any encryption. This reflects an optimistic assessment of these approaches given that using the trained models under HE to generate predictions may incur an additional loss of precision. Note that HE frameworks are based on modular integer arithmetic and thus do not naturally support high-precision floating-point operations. For both HE-based methods, we downloaded the software published by the authors (https://github.com/K-miran/secure-imputation) and evaluated the methods on our datasets using the suggested default parameters. Due to the computational burden of training a predictive model for every test variant, we sampled 1000 test variants uniformly at random for each MAF category to approximate the *r*^2^ value for each subject. Our results for Minimac and SMac are also based on the same set of test variants for the fairness of comparison. Both on 1KG and HRC datasets, linear models used by HE-based solutions produced significantly lower imputation accuracies in all three MAF categories, with HE-EPFL consistently obtaining more accurate results than HE-UTMSR (Figure 3). For example, test variants with MAF between 0.5 and 5% were imputed with average *r*^2^ value of 0.88 for Minimac/SMac, 0.70 for HE-EPFL, and 0.58 for HE-UTMSR. Inspecting the *r*^2^ values of the individual test subjects revealed that the gap in accuracy between Minimac/SMac and the HE-based solutions are consistent across nearly all subjects. These observations are in line with the intuition that HMM-based imputation methods such as Minimac can leverage more complex and long-range correlation patterns in the genotype dataset than what locally trained linear models in the HE-based solutions can capture. Overall, our results show that SMac is a compelling solution for secure genotype imputation that provides state-of-the-art accuracy along with our strengthened leakage-resilience under the SGX model.

### SMac efficiently scales to large datasets in both runtime and memory usage

Deploying a software to run inside an SGX enclave could introduce a performance bottleneck due to the additional computational overhead in encrypting and decrypting the data stored in the enclave memory as well as the limited amount of memory available to an SGX process. This raises the potential concern that SGX-based secure genotype imputation may incur overwhelming computational overhead for the emerging large-scale genetic datasets (e.g. the TOPMed reference panel, which is approaching 100k individuals [51]). To address this issue, we evaluated the scalability of SMac both in terms of runtime and memory usage for a range of datasets of varying sizes, including 1KG (~ 5k haplotypes), uniformly downsampled HRC datasets (10k and 25k haplotypes), and the full HRC dataset (~ 54k haplotypes). For completeness, we additionally tested a variant of SMac, termed SMac-lite, which corresponds to an SGX-based solution that implements the Minimac algorithm in the same way as SMac, but without the additional protection mechanisms we introduced for side-channel leakages. We measured the performance metrics for imputing the entire chromosome 20, which is performed by sequentially processing each of the three overlapping segments of the chromosome generated by Minimac in the case of SMac/SMac-lite. All our experiments were run on a system with an Intel SGX enabled Intel Xeon E-2288G processor, with 16 GB RAM and 112 MB enclave page cache (EPC; i.e. amount of memory available to an SGX process). Memory usage that exceeds the EPC limit is swapped to RAM with authenticated encryption of memory pages, relying on the native feature of SGX. Data transfer between the client and the server was simulated via loopback TCP and peak heap usage measurements were taken with massif (valgrind), a standard memory profiling tool for Linux.

Our experiments demonstrated a linear scaling of both runtime and memory with respect to reference panel size for all of the methods evaluated (Figure 4). As expected, SMac shows an overhead in runtime (54% additional runtime on the full HRC dataset), but we believe that this additional burden is a small price compared to the value of enhanced privacy protection that our solutions additionally provide to the users. Note that under severe runtime constraints, SMac-lite may be a viable alternative, considering its fast runtime, which is 36% lower on average than that of Minimac, owing to our efficient implementation with respect to memory usage and utilization of native CPU features. Although SMac-lite does not benefit from our protection mechanisms against side-channel leakages and thus remains vulnerable to sophisticated attackers with sufficient incentives, we do note that SMac-lite still runs the imputation pipeline entirely inside the SGX enclave, thus proving a stronger security guarantee than the status quo of uploading the user’s raw data to a third-party server.

**Figure 4:**
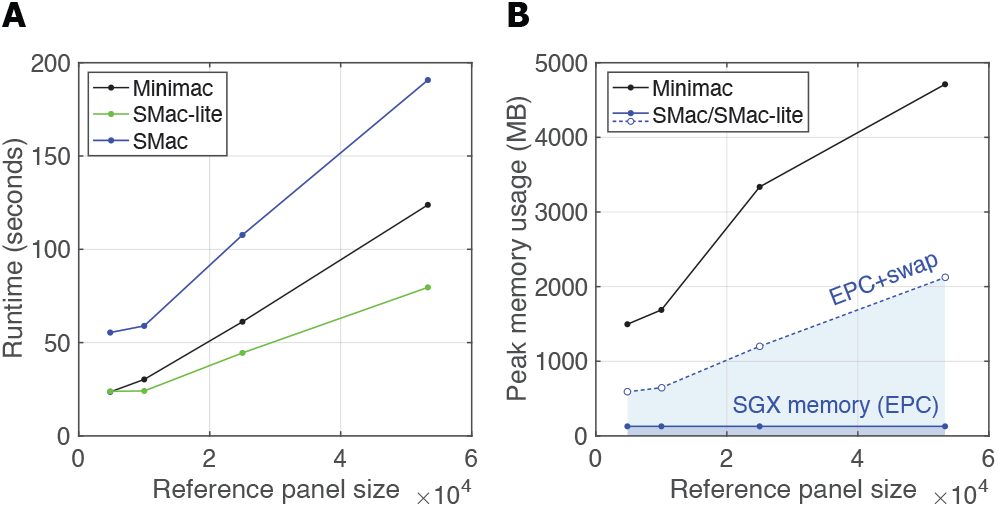
SMac achieves practical performance with respect to runtime and memory usage. We measured the runtime (**A**) and memory usage (**B**) of SMac and Minimac4 for imputing chromosome 20 of a single sample on a range of reference panels with varying sizes, including 1KG, HRC, and subsampled HRC datasets with 10k and 25k haplotypes each. We also show the performance of SMac-lite, a less-secure alternative to SMac which runs the same imputation algorithm as SMac in SGX without our additional protection for side-channel leakages. All results reflect the average of five trials. All methods show linear scaling in both runtime and memory with respect to data size, and SMac incurs a modest 54% runtime overhead on the largest dataset with 54k haplotypes while additionally providing protection for the user’s data. SMac and SMac-lite use *~*100 MB in enclave page cache (EPC) and the rest in non-EPC RAM for swap (total is shown as EPC+swap). This overall halves the memory usage of Minimac4.

With respect to memory usage, SMac and SMac-lite are identical, and they both use significantly less memory than Minimac (e.g. 55% smaller on full HRC data). This is due to the optimized usage of memory in our algorithmic implementation, additionally aided by Rust’s effective memory management techniques. Recall that any excess memory allocation beyond the EPC limit is swapped to RAM, which adds to the computational overhead of SGX applications; however, SMac/SMac-lite has a simple memory access pattern due to the sequential nature of the imputation algorithm, which alleviates the burden of swap memory usage as indicated by the modest runtime overhead of our tools. These results show that our SGX-based approach to secure genotype imputation is nearly as practical as existing imputation pipelines and will remain applicable for emerging large-scale reference datasets. Broadly, our work provides a useful methodological guideline for how state-of-the-art algorithms and software tools for analyzing sensitive genomic datasets could be ported to the trusted execution environments in a secure and efficient manner.

## Acknowledgements

We thank Hongbo Chen and Weijie Liu for their comments on the known attack surfaces of SGX technology and mitigation strategies.

## Footnotes

† This manuscript has been accepted for oral presentation at RECOMB 2021.

