## Supplementary Information for "Privacy-Preserving Genotype Imputation in a Trusted Execution Environment"

### Appendix

#### A. State Space Reduction of SMac and Minimac

In this section we give a detailed review of the "state space reduction" approach employed by SMac, as per Minimac (in particular Minimac3 [11]). We start by the problem definition and notations used in the description of our algorithm. Then we describe the commonly used Hidden Markov Model (HMM) for genotype imputation as well as our approach to reduce the number of hidden states in the HMM.

**HMM-based genotype imputation.** We consider the genotype imputation problem as follows: Given a *target haplotype* with a sequence of $n$ genotypes $S = (S_1, \cdots, S_n) \in \mathcal{A}^n$, some of which are missing (to be imputed), where $\mathcal{A}$ represents the alphabet (e.g. $\mathcal{A} = \{A, C, G, T\}$); and (ii) a *reference panel*, which consists of $m$ fully-observable haplotypes $R^{(1)}, \cdots, R^{(m)} \in \mathcal{A}^n$. Let $O \subset [n]$ be the subset of indices where $S_i \in O$ is observed in the target haplotype. Let $M = [n] - O$ be the set of missing genotypes in the target haplotype to be imputed. The goal is to infer the missing genotype $S_i$ for each $i \in M$, based on the input reference panel.

At a high level, the HMM used in SMac models $S$ as a *mosaic* of haplotypes in the reference panel. Specifically, we assign a random variable $X_i \in [m]$ for each $S_i$ to indicate (at genotype $i$) which reference haplotype the target haplotype shares the genotype with. We refer to $[m]$ the *state space* of our HMM. Let $S_I = (S_{i_1}, S_{i_2}, \cdots, S_{i_{|I|}})$ for $i_1 < i_2 < \cdots < i_{|I|} \in I \subset [n]$ denote the sequence of genotypes on a subset $I$ of the $n$ positions (which can be defined similarly on each $R^{(j)}$). Then we want to compute for each $i \in M$, $\mathbb{P}(S_i | S_O, \theta)$, where $\theta = (T_i(j,l), E_j(S_i), \pi_j)$ corresponds to the parameters in the HMM, including

- Transition probabilities (for $i = 1, \cdots, n-1$, $j, l = 1, \cdots, m$)

$$T_i(j,l) = \mathbb{P}(X_{i+1} = l | X_i = j) = \begin{cases} 1 - \lambda_i + \dfrac{\lambda_i}{m} & \text{if } l = j \\ \dfrac{\lambda_i}{m} & \text{if } l \neq j \end{cases}$$

  where $\lambda_i$ denotes the recombination rate between haplotypes at position $i$.

- Emission probabilities (for $i = 1, \cdots, n$, $j = 1, \cdots, m$ and $S_i \in \mathcal{A}$)

$$E_j(S_i) = \mathbb{P}(S_i | X_i = j) = \begin{cases} \epsilon & \text{if } i \in O \text{ and } S_i \neq R_i^{(j)} \\ 1 - \epsilon & \text{if } i \in O \text{ and } S_i = R_i^{(j)} \\ 1 & \text{if } i \notin O \end{cases}$$

  where $\epsilon$ denotes the probability of genotyping error, i.e., the underlying and observed genotypes do not match.

- Initial probabilities $\pi_j = \mathbb{P}(X_1 = j)$ (for $j = 1, \cdots, m$), which can be set to uniform distribution, i.e., $\mathbb{P}(X_1 = j) = \frac{1}{m}$ for $j = 1, \cdots, m$.

We drop $\theta$ in the following expressions for simplicity. Note that it suffices to compute each $\mathbb{P}(X_i = j | S_O)$ for $j \in [m]$, since

$$\mathbb{P}(S_i | S_O) = \sum_{j=1}^{m} \mathbb{P}(S_i | X_i = j) \mathbb{P}(X_i = j | S_O);$$

while $\mathbb{P}(X_i = j | S_O)$ can be computed using the well-known forward-backward algorithm [28]. Let $F_i(j)$ and $B_i(j)$ denote the forward and backward probabilities for each state $j \in [m]$ at genotype $i$ respectively, then we can update respectively

$$F_{i+1}(j) = \left[ (1 - \lambda_i) F_i(j) + \frac{\lambda_i}{m} \sum_{l=1}^{m} F_i(l) \right] \cdot E_j(S_{i+1})$$

for $i = 1, \cdots, n-1$ ($F_1(j) = \pi_j \cdot E_j(S_1)$) and

$$B_{i-1}(j) = \frac{\lambda_{i-1}}{m}\left(\sum_{l=1}^{m} B_i(l)E_l(S_i)\right) + (1-\lambda_{i-1})B_i(j)E_j(S_i)$$

for $i = n-1, \cdots, 2$ ($B_{n-1}(j) = \frac{\lambda_{n-1}}{m}\sum_{l=1}^{m} E_l(S_i) + (1-\lambda_{n-1})E_j(S_i)$). Finally, the posterior $\mathbb{P}(X_i = j|S_O)$ is given by

$$\mathbb{P}(X_i = j|S_O) \propto F_i(j) \cdot B_i(j).$$

**State space reduction.** The key step of the state space reduction approach, introduced in Minimac3 [11], is to partition the genotypes of the target haplotype into $K$ consecutive *blocks* $c_1, \cdots, c_K$, where each pair of consecutive blocks $c_k$ and $c_{k+1}$ overlap at one position $i_k$. Consider a specific block $c_k = (i_{k-1}, i_{k-1}+1, \cdots, i_k)$ for $k \in [K]$. Let $\mathcal{S} = [m]$ be the original state space. Let $\mathcal{S}' = \{R_{c_k}^{(j)} \mid j \in [m]\} = \{R_{c_k}^{(1)}, \cdots, R_{c_k}^{(m_k)}\}$ (without loss of generality we can assume $\mathcal{S}' = [m_k]$) be the *reduced* state space, i.e., the collection of distinct haplotypes within block $c_k$; and $Y_i \in [m_k]$ be a random variable indicating the corresponding state at position $i$. Let $m_{j'} = \sum_{j=1}^{m} I(R_{c_k}^{(j)} = R_{c_k}^{(j')})$ for $j' \in [m_k]$ be the number of haplotypes in the reference panel that matches $R_{c_k}^{(j')}$ within block $c_k$, where $I(\cdot)$ is the indicator function. In this case we only need to update within each block the forward and backward probabilities defined on the reduced state space $\mathcal{S}'$, i.e. for each $j' \in [m_k]$:

$$F_{i+1}(j') = \left[(1-\lambda_i)F_i(j') + \frac{m_{j'}\lambda_i}{m}\sum_{l'=1}^{m_k} F_i(l')\right] \cdot E_{j'}(S_{i+1})$$

for $i = i_{k-1}, \cdots, i_k - 1$;

$$B_{i-1}(j') = \frac{m_{j'}\lambda_{i-1}}{m}\left(\sum_{l'=1}^{m_k} B_i(l')E_{l'}(S_i)\right) + (1-\lambda_{i-1})B_i(j')E_{j'}(S_i)$$

for $i = i_k, \cdots, i_{k-1}+1$. We note that the emission probabilities at genotype $i = i_{k-1}, \cdots, i_k$ should agree among those $j \in [m]$ with $R_{c_k}^{(j)} = R_{c_k}^{(j')}$ - therefore, we denote the emission probabilities as $E_{j'}(S_i) = \mathbb{P}(S_i|Y_i = j')$ in the reduced state space.

Additionally, we need to transform the forward and backward probabilities defined on the reduced state space to those defined on the original state space at block boundaries $i_{k-1}$ and $i_k$.

$$F_{i_{k-1}}(j') = \sum_{j=1}^{m} I(R_{c_k}^{(j')} = R_{c_k}^{(j)}) \cdot F_{i_{k-1}}(j)$$

$$F_{i_k}(j) = \frac{1}{m_{j'}} \cdot F_{i_k}^{R}(j') + \frac{F_{i_{k-1}}(j)}{F_{i_{k-1}}(j')} \cdot F_{i_k}^{NR}(j')$$

$$B_{i_k}(j') = \sum_{j=1}^{m} I(R_{c_k}^{(j')} = R_{c_k}^{(j)}) \cdot B_{i_k}(j)$$

$$B_{i_{k-1}}(j) = \frac{1}{m_{j'}} \cdot B_{i_{k-1}}^{R}(j') + \frac{B_{i_k}(j)}{B_{i_k}(j')} \cdot B_{i_{k-1}}^{NR}(j')$$

where $F_i^R(j')$ and $F_i^{NR}(j')$ ($i_{k-1} \leq i \leq i_k$) denote the component of forward probabilities between genotypes $i_{k-1}$ and $i$ without recombination, or with at least one recombination event respectively; $B_i^R(j')$ and $B_i^{NR}(j')$

are defined similarly for backward probabilities. These probabilities can be updated as:

$$F_i^{NR}(j') = F_{i_{k-1}}(j') \prod_{i'=i_{k-1}}^{i-1} \left[(1-\lambda_{i'}) \cdot E_{j'}(S_{i'+1})\right]$$
$$F_i^{R}(j') = F_i(j') - F_i^{NR}(j')$$
$$B_i^{NR}(j') = B_{i_k}(j') \prod_{i'=i}^{i_k-1} \left[(1-\lambda_{i'}) \cdot E_{j'}(S_{i'+1})\right]$$
$$B_i^{R}(j') = B_i(j') - B_i^{NR}(j')$$

Finally, the posterior probability $\mathbb{P}(Y_i = j'|S_O)$ is given by

$$\mathbb{P}(Y_i = j'|S_O) \propto \left(\sum_{j=1}^{m} I(R_{c_k}^{(j')} = R_{c_k}^{(j)}) F_{i_{k-1}}(j) B_{i_k}(j)\right) \cdot \left(\frac{F_i^{NR}(j')}{F_{i_{k-1}}(j')} \cdot \frac{B_i^{NR}(j')}{B_{i_k}(j')}\right) + \frac{1}{m_{j'}}\left(F_i(j')B_i(j') - F_i^{NR}(j')B_i^{NR}(j')\right)$$

and $\mathbb{P}(S_i|S_O)$ is given by

$$\mathbb{P}(S_i|S_O) = \sum_{j'=1}^{m_k} E_{j'}(S_i)\mathbb{P}(Y_i = j'|S_O).$$

It was shown in [26] that using the reduced state space as "templates" will produce exactly the same imputation result $\mathbb{P}(S_i|S_O)$ as that is computed by using the original state space, i.e.,

$$\sum_{j'=1}^{m_k} E_{j'}(S_i)\mathbb{P}(Y_i = j'|S_O) = \sum_{j=1}^{m} E_j(S_i)\mathbb{P}(X_i = j|S_O).$$

However, the running time of the forward-backward recursion in the reduced state space is $\Theta((\sum_{k=1}^{K} m_k \ell_k) + Km)$ (where $\ell_k = i_k - i_{k-1} + 1$ is the number of genotypes in block $c_k$), which, in practice, is much smaller than $\Theta(mn)$ to run the basic forward-backward recursion in the original state space, since usually we have $m_k \ll m$. The optimal partition (which minimizes $\sum_{k=1}^{K} m_k \ell_k + \Theta(Km)$) can be found through dynamic programming, as introduced in [11].

SMac takes advantage of the same reduced state space generated by Minimac for each block in the optimal partition. More specifically, we assume that the reference panel, available to the imputation service provider in plaintext, is preprocessed with Minimac3 to generate the M3VCF file (provided as input to SMac) which compactly represents the panel in the reduced state space. In addition, we consider the HMM parameters as given to SMac - which means that the parameter estimation step does not need to be performed securely in SGX - since they are estimated from the reference panel. [‡]

#### B. Leakage-resilient approximation of Log-Sum-Exp (LSE) and Log-Diff-Exp (LDE) functions

Suppose that $x = \ln(a)$ and $y = \ln(b)$, then

$$\ln(a+b) = \ln\left(e^{\ln(a)} + e^{\ln(b)}\right) = \ln\left(e^x + e^y\right) \approx \max(x,y) + \ln(1 + e^{-|x-y|}) \quad (1)$$

$$\text{given } a > b,\ \ \ln(a-b) = \ln\left(e^{\ln(a)} - e^{\ln(b)}\right) = \ln\left(e^x - e^y\right) \approx x + \ln(1 - e^{-|x-y|}) \quad (2)$$

[‡]Minimac estimates the recombination and error rates by maximizing the likelihood for a given reference panel based on a combination of the expectation-maximization (EM) and Monte Carlo Markov chain (MCMC) sampling algorithms [11].

We define $\texttt{LSE}(x, y) := \max(x, y) + \ln(1 + e^{-|x-y|})$ and $\texttt{LDE}(x, y) := x + \ln(1 - e^{-|x-y|})$. In order to compute the expressions equivalent to $a + b$ and $a - b$ in log space, we need to approximate LSE and LDE where $x$ and $y$ are timing-protected log-transform fixed points. Notice that the term $\max(x, y)$ in LSE can be easily computed with a simple boolean circuit without side-channel leakage. Our challenge then is to approximate $\ln(1 + e^{-|x-y|})$ and $\ln(1 - e^{-|x-y|})$ (given $x > y$).

We define $\texttt{NLS}(a) := \ln(1+e^{-a})$ and $\texttt{ODE}(a) := \ln(1-e^{-a})$ and observe that $\texttt{NLS}(a)$ and $\texttt{ODE}(a)$ converges to zero quickly with respect to $a$. This makes the two functions amenable to polynomial approximation because after some small $a > 15$, we can approximate $\texttt{NLS}(a) = \texttt{ODE}(a) \approx 0$. Recall that timing-protected polynomial approximation is practical because addition, subtraction, and multiplication of integers are constant-time instructions. For practical efficiency, we choose 4-piecewise polynomial approximation of degree 2 where the effective range of $a$ ($0 < a < 15$) is split into 4 equal sections. This approximation technique gives us reasonable accuracy without incurring too much computational cost. Our experiment in Section 3 demonstrates that SMac's performance under this technique in SGX is competitive with Minimac.

#### C. Strong Timing-Protected Typing

We formally define the typing rules of SMac program, which systematically ensure timing protection of the overall program *at the syntactic level*. Intuitively, the typing rules enforce the consistency of timing protection; once a value becomes timing-protected (i.e. only allowed to be passed into routines that has no timing dependence on the input), any further processing of the value remains timing-protected. Unsafe operation over a value is allowed only if it is explicitly exposed.

We first identify the types and their associated binary and unary operations required for SMac implementation: user's genotypes, booleans, and integers. We have verified that these types and operations are sufficient for SMac's correctness and security.

$$\texttt{geno} := \texttt{Reference} \mid \texttt{Alternative} \mid \texttt{Missing} \quad (3) \qquad \texttt{v} := \texttt{geno} \mid \texttt{bool} \mid \texttt{int} \quad (4)$$

$$* := \texttt{and} \mid \texttt{or} \mid \texttt{xor} \mid < \mid > \mid = \mid \texttt{lshift} \mid \texttt{rshift} \mid + \mid - \mid \times \quad (5) \qquad \hat{*} := \neg \mid - \quad (6)$$

Notice the lack of $\div$. This is because integer division is not constant-time. We have shown how to get around this issue in Section 2.4. A value can be marked timing-protected via operation $\texttt{protect}\,(\cdot)$ and unmarked via $\texttt{expose}\,(\cdot)$.

$$x : \texttt{v} \ : \ \texttt{protect}\,(x) \to \texttt{Tp}\,[x] \quad (7) \qquad x : \texttt{v} \ : \ \texttt{expose}\,(\texttt{Tp}\,[x]) \to x \quad (8)$$

Once an unmarked value interacts with a marked value, the outcome is always marked. A marked value always stay marked.

$$\begin{aligned} a : \texttt{v}, b : \texttt{v}, c : \texttt{v} \ : \ a * b \to c \\ \implies a * \texttt{Tp}\,[b] \to \texttt{Tp}\,[c] \ \wedge \\ \texttt{Tp}\,[a] * b \to \texttt{Tp}\,[c] \ \wedge \\ \texttt{Tp}\,[a] * \texttt{Tp}\,[b] \to \texttt{Tp}\,[c] \end{aligned} \quad (9)$$

$$a : \texttt{v}, b : \texttt{v} \ : \hat{*} a \to b \implies \hat{*}\texttt{Tp}\,[a] \to \texttt{Tp}\,[b] \quad (10)$$

Finally, a timing-protected boolean can be used for branching via operation $\texttt{select}\,(\cdot)$. Any values produced by the branching become timing-protected.

$$\begin{aligned} x : \texttt{bool}, a : \texttt{v}, b : \texttt{v}, c : \texttt{v} \ : \ \{\texttt{if } x \texttt{ then } c = a; \texttt{else } c = b\} \implies & \texttt{select}\,(\texttt{Tp}\,[x]\,, a, b) \to \texttt{Tp}\,[c] \ \wedge \\ & \texttt{select}\,(\texttt{Tp}\,[x]\,, a, \texttt{Tp}\,[b]) \to \texttt{Tp}\,[c] \ \wedge \\ & \texttt{select}\,(\texttt{Tp}\,[x]\,, \texttt{Tp}\,[a]\,, b) \to \texttt{Tp}\,[c] \ \wedge \\ & \texttt{select}\,(\texttt{Tp}\,[x]\,, \texttt{Tp}\,[a]\,, \texttt{Tp}\,[b]) \to \texttt{Tp}\,[c] \end{aligned} \quad (11)$$

#### D. Other Implementation Details

In this section, we provide addition implementation details according to the mitigation strategies not mentioned in the main paper.

**SGX Remote Attestation in SMac** The SGX remote attestation workflow is visualized in the right panel of Figure 1. The remote attestation protocol requires that the Service Provider is deploying SMac inside an SGX enclave on a SGX-capable device, whereas User can interact with the server from any device running the remote attestation software. The protocol proceeds as follows.

1. Service Provider initializes SMac with the reference panel inside an SGX enclave.
2. Service Provider generates a report of the running SMac binary (*without* the reference panel) via the Architectural Enclave Service Manager (AESM). This report generally contains a hash of the SMac binary, SGX firmware information, and the first half of Diffie-Hellman key exchange, all of which signed with the SGX hardware's secret signing key. The report is sent to User.
3. User passes the report to the Intel Attestation Service (IAS) for signature verification. If the report is valid, the SMac binary hash is verified for authenticity (i.e. that the source code has not been tampered with). Lastly, User generates the second half of Diffie-Hellman key exchange and sends it to Service Provider.
4. User and SGX enclave on Service Provider's side establish a secure channel with the shared Diffie-Hellman key. User sends genotypes to the enclave via the secure channel. Once the genotypes are imputed, the enclave returns them to User via the secure channel.

**Rust Programming Language.** Rust is advantageous for SMac implementation and therefore is our choice for a number of reasons. *First*, implementing the typing rules described in Appendix C is native to Rust: SMac simply defines a new `struct` as a zero-cost wrapper around existing integer types and overrides all arithmetic operations with their leakage-resilient log-transformed version. *Second*, Rust's memory model helps prevent memory bugs as well as reduce memory usage by deallocating memory as soon as it becomes obsolete. SMac's low memory usage in comparison with Minimac is largely thanks to this feature. *Third*, Rust is a low-level language; this grants SMac complete control over security-critical code on an assembly level. And *fourth*, Rust's user-friendly package manager `cargo` and crate registry `https://crates.io` allow us to take advantage of many existing efficient libraries without all the hassle of dependency management.

**Native CPU optimization.** We utilize Intel's `AVX` and `SSE` features for efficient single-instruction-multiple-data (SIMD) integer arithmetics. This is done through Rust's built-in LLVM loop-vectorization and our effective use of the `rust-ndarray` library for linear algebra.

**Timing Protection.** Implementation leakage-resilient operations in Section 2.4 in practice is challenging because modern compilers are equipped with techniques to optimize around artificial barriers introduced by programmers to enforce timing protection. For example, if `select` $(\cdot)$ is implemented for timing protection in a straight-forward manner, Rust compiler is likely to know how to take a shortcut without needing to compute both branches. To overcome this challenge, Andrysco *et al.* [20] manually inspect the assembly code of their `fixed-time-fixed-point` library to ensure expected behaviors, and suggest that users link the library binary as provided without recompilation. We find approach unsustainable for good performance because it also prevents all desirable compiler optimizations.

Instead, we follow `rust-timing-shield` library's strategies [53] in two ways. *First*, because Rust compiler will *not* optimize in-line assembly, in-line assembly can be used to create predictable behaviors. The drawback of this strategy is likewise the lack of optimization which can impact the overall performance, although the impact is far less than linking a whole library this is pre-compiled. *Second*, because problematic Rust compiler optimization always occurs at boolean operations, we can prevent it by "tricking" the compiler into misrecognizing protected booleans by inserting an empty in-line assembly block. SMac utilizes both strategies to balance between security and performance.

**Fortanix EDP.** Fortanix EDP is a Intel SGX framework for Rust. We choose this framework over the alternative because it is designed for network services to make development and deployment simple. Fortanix EDP is also compatible with most existing Rust code; this is indeed an advantage over Intel SGX SDK in `C++` because existing libraries can be simply plugged in without manual porting. One caveat is Fortanix EDP is still in early development which requires Nightly Rust to compile.
